# Adapting Social Operant Paradigms to Measure Postpartum Maternal Motivation

**DOI:** 10.64898/2026.08.20.746000

**Authors:** Sydney Ku, Judy Nyakoa, Gonzalo Miranda, Debra A. Bangasser

## Abstract

Operant paradigms are powerful tools to quantify motivation and reward. Traditionally, operant conditioning research has been limited to food and drug reinforcers. Recent advances in commercially available operant equipment, however, allow for the quantification of social motivation. These operant assays are an improvement over commonly used social preference tasks, as they enable direct measurement of the effort and motivation driving social behavior. Based on a design by Venniro et al. (2020), the MedPC social operant boxes modify the traditional operant box setup for social interactions. The experimental rat can lever-press to raise a door for an interaction with a target rat behind a porous barrier. These social operant boxes have been widely adapted to test social behavior in adult and adolescent rodents and investigate how a range of conditions (e.g. stress, drug taking, etc.) affect social motivation. However, there is a gap in assessing maternal motivation for pups during the postpartum period, despite ample evidence that postpartum social behavior is highly relevant for offspring health outcomes. Here, we detail 3D-printed modifications to the standard Med PC social operant boxes to adapt the social target chamber to safely house neonatal pups. We have also developed testing protocols to assess motivation during the limited postpartum period. These data demonstrate that, with simple modifications to social operant chambers and testing protocols, the field can implement advanced behavioral approaches to directly assess maternal motivation.

## INTRODUCTION

The importance of maternal behavior, specifically care, in offspring health is well-established.^1^ Early life experiences have a profound role in shaping developmental trajectories and behavioral phenotypes^2–5^. The quantity and frequency of parental care have been shown to scaffold brain development, behavior, and health for offspring across taxa^2,6–8^. In humans, poor parental caregiving or neglect are associated with higher risk for neuropsychiatric diseases in offspring^9,10^. Maternal neglect, for instance, is associated with increased risk of substance abuse, attention-deficit/hyperactivity disorder, depression, anxiety, and bipolar disorders in adulthood^11–13^. Given the known effects of maternal caregiving, examining the neurobiological underpinnings for maternal behaviors and offspring motivation is critical.

Basic neuroscience approaches and animal models allow for control of environmental factors, allowing elucidation of the effect of experience on behavioral phenotypes. Preclinical rodent research has delineated compelling evidence for stable, naturally occurring individual differences in maternal care behaviors^2,3,14–16^. Maternal care is considered a passive or active behavior on behalf of a mother directed toward the development (nursing, licking, grooming, etc.) or protection (nesting, defense, retrieval) of her young. In rodents, the most widely adopted approach to understanding maternal motivation involves recording care behaviors, particularly pup-directed licking, grooming, carrying, nursing, or nestbuilding^4,7,14,17^.

Naturally occurring variations in dam (a rat mother) care have been proven to predict individual differences in later-life offspring health outcomes^14,15^. Pups exposed to low levels of maternal care, for example, show altered hypothalamic-pituitary-adrenal (HPA) axis functioning, gene expression, and dopaminergic activity^3,15,16,18,19^. Behaviorally, pups receiving low care consume more alcohol and cocaine and show increased anxiety and depressive-like behaviors, analogous to the elevated human risk for substance abuse and mood disorders associated with maternal neglect^20,21^. Interestingly, pups receiving high levels of care exhibit more modest HPA axis responses to stress and show fewer behavioral signatures of fear^22^. Cross-fostering has validated that maternal care, rather than genetics, are responsible for these shifts in behavior and stress-responsivity^22^. Interestingly, exposure to high or low maternal care phenotypes have distinct epigenetic markers, such as levels of stress hormones, expression of stress receptors, and neurotransmittic markers^23^. These studies point to a clear role for maternal attention and behavior in offspring health and fitness. They underscore the importance of early life tactile experiences in developing stress responsivity, while showing that maternal care, specifically, can promote resilience or risk for poor outcomes. While care remains an important and relevant assortment of behaviors, investigating mechanisms underlying maternal motivation is crucial, yet understudied.

Postpartum social assays have been developed to test maternal behaviors towards pups. Pup retrieval paradigms, for example, involve scattering pups across the corners of the dam’s home cage and measuring the amount of time it takes for her to bring them back to the nest^24–26^. Retrieval assays mimic a common form of maternal responsivity warranted in the wild, representing an ethologically valid approach to quantifying motivation^27^. Similarly, conditioned place preference (CPP) assays can be used to compare motivation for competing reinforcers and have been effectively applied to postpartum behavior^28,29^. These studies have been integral in comparing the rewarding value of reinforcers and identifying key brain regions for pup-directed motivation, such as the medial preoptic area (mPOA)^30–34^. Examining naturalistic maternal behaviors (modelled with pup retrieval or maternal care) or the associative power of reinforcers (modelled with CPP) has been critical to understanding maternal motivation.

While these assays are effective, operant approaches represent an advantage to quantifying motivation, as they model volitional aspects of the reward. Operant conditioning allows for the study of a wide range of variables, such as the ratio of responses, the time interval between strongest responses, and variations in stimuli that signaled the opportunity to earn rewards^35^. Most critically, the operant apparatus quantifies the response rate – the measure of response strength across time. While the response rate is a continuous variable, it is sensitive to environmental perturbations, stress manipulations, and subject health, and thus represents a reliable way of quantifying reinforcer motivation across experimental conditions and time points^35,35–37^. Importantly, operant conditioning boxes model volitional taking of a reinforcer, as opposed to simply quantifying associations^38–41^. Further, examining motivation as a function of a behavioral consequence, such as lever pressing, is highly translatable, much akin to human taking of drugs, acquisition of food, or reaching out for social support.

Given these operant advantages, it is not surprising that two groups used modified operant chambers to investigate maternal motivation for pups^42,43^. Lee et al., (1999) trained postpartum rats to press a lever for an interaction with a pup, which a researcher would manually administer through a shoot. Hauser and Gandelman (1985) built a testing apparatus where the home cage could be inserted into a detachable operant chamber, and trained mice to press lever-press for their pups. While these methods were useful in understanding maternal motivation, they were not widely adopted. This is likely because the method required researcher intervention (i.e., it was not automated) and the equipment was not commercially available. Since this publication, Venniro et al. (2020), developed an automated social operant box design (made commercially accessible by MedAssociates) where a rodent can press a lever for a social interaction with a conspecific^39,44^. Specifically, when the lever is pressed by the experimental rat, a guillotine door rises, revealing a target rat behind a porous barrier. The barrier allows for tactile, olfactory, and auditory cues, but keeps the rats separate so as not to require researcher intervention (i.e., there is no need to return rodents to their respective chambers after each interaction). These social boxes are equipped with food and drug reinforcers, allowing for choice studies where rats can choose between food, drug, or social rewards.^39^ These chambers have been quickly adopted and used to reveal how a range of stimuli, from stressors to development, affect rodent social motivation^36^.

Despite their widespread use, these social boxes have not yet been implemented to assess maternal motivation due to several challenges. First, the target social chamber is not designed to safely house and contain pups. Second, the window to test maternal motivation is brief; the earliest pups can be put in the operant chamber is when they can thermoregulate on PND11^45–47^. Yet maternal care and motivation wane around PND16, when pups develop executive motor skills and more sophisticated sensory systems^17^. Typical operant training schedules exceed this short window. To address these obstacles, we 1) 3D-printed inserts to securely and comfortably contain pups in the social target chamber (Fig. 1); and 2) adjusted testing parameters to accommodate the short window for maternal motivation and pup thermoregulation (Fig. 2). To abbreviate the postpartum operant assessment, we leveraged the chamber’s versatility for sucrose self-administration and trained the females to lever press as virgins. Prior to pregnancy, they learned to press a separate lever to obtain sucrose pellets. The virgin sucrose training ensured that rats understood the lever-reward contingency and were habituated to the chambers prior to the postpartum assessment. We then were able to truncate postpartum testing to fewer days (1 day of each fixed ratio training) with shorter assessments (30 minutes each) to accommodate the brief window of testable motivation. Using this method, we developed a replicable and highly accessible operant protocol to test postpartum social motivation for pups and elucidated behavioral profiles for pup-directed maternal motivation.

**Figure 1.**
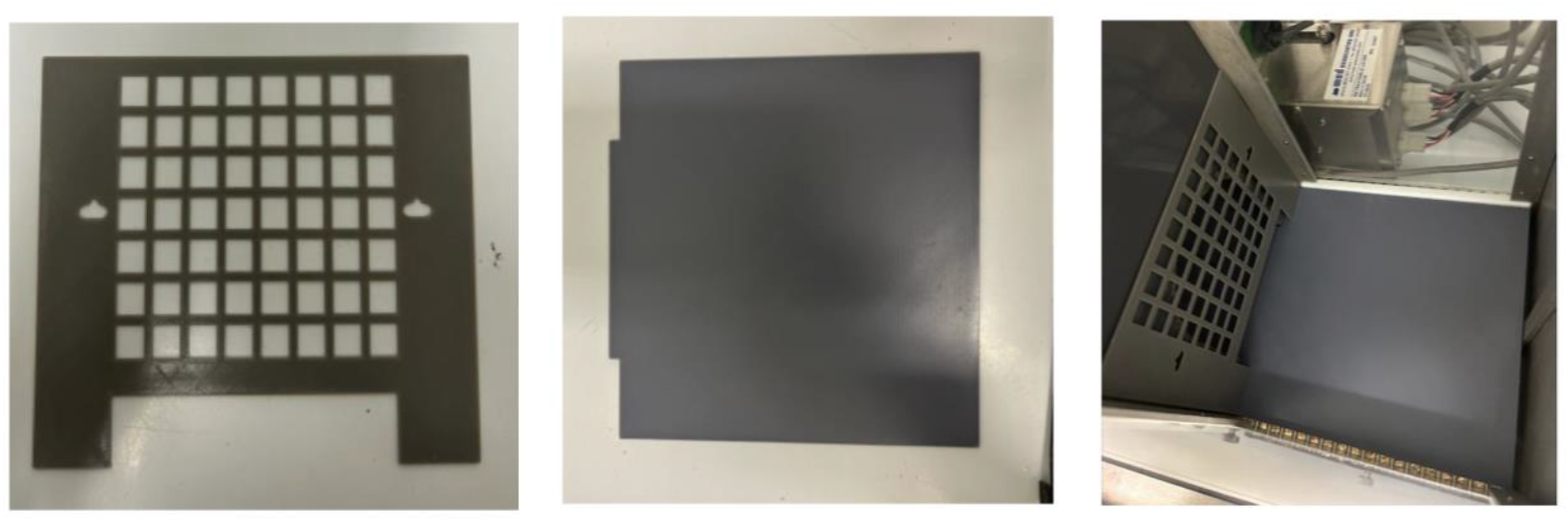
Modifications to Med Associates Social Operant Box. We modified aspects of the standard rat social operant Med Associates chambers to safely house pups using 3D printed parts. **A) Social door attachment.** Modified social door has a smaller grid size than the standard door to account for pup size. **B) Social operant chamber floor cover.** Floor cover for the social chamber to prevent pup stress from standard rod flooring. **C) Affixed modifications.** Example of the social door attachment and social operant chamber floor cover as needed for behavior.

**Figure 2.**
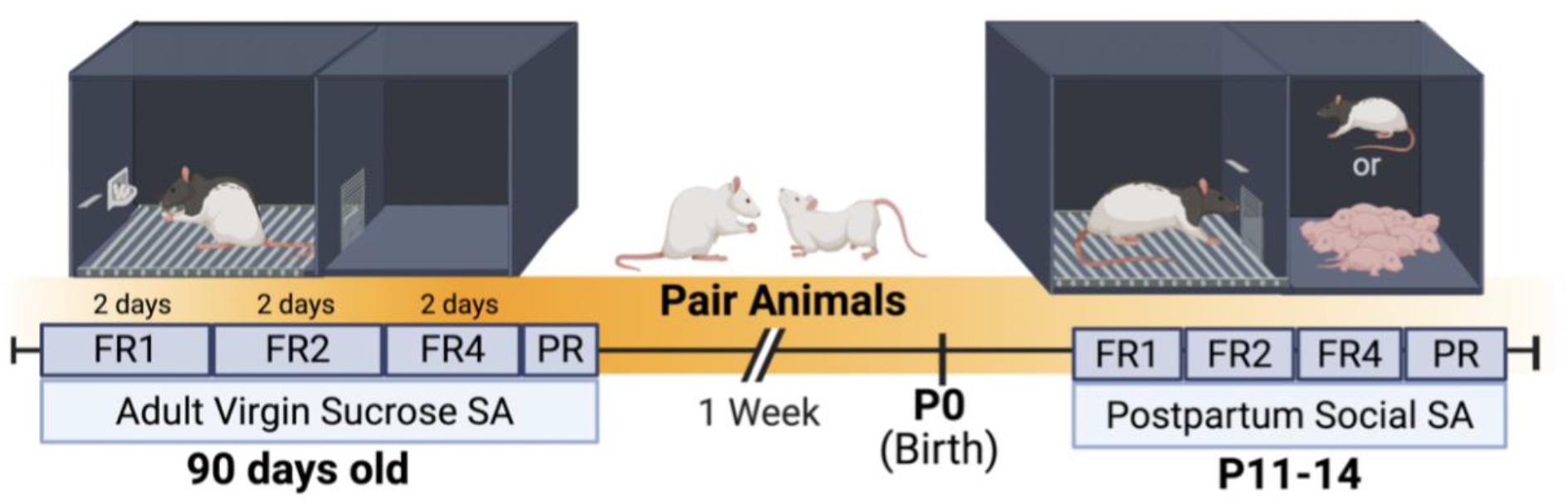
Experimental Timeline. Before pregnancy, adult female Long-Evans rats were trained to press a lever for sucrose pellets to acquire the **self-administration (SA)** lever-reward paradigm. After completing sucrose SA training, females were paired for breeding. On PND11, postpartum dams were tested in a social operant SA task, where pressing a designated social lever granted access to either their pups or an adult female conspecific. Tissue was collected an hour after the final behavioral assessment.

## METHODS

### MedAssociates Social Boxes

Venniro et al. (2020), provided a standardized protocol for social operant self-administration to quantify social reward in rats using MedAssociates boxes ^44^. This protocol included discrete-trial choices between a social interaction and additional reinforcers, such as drugs or sucrose, to compare motivational properties. These social boxes provide a reliable way of studying the role of social reward in rodent models, as well as vulnerability to motivational dysregulation. The standard MedAssociates modular operant test chamber for rats (11.63” × 9.78” ×7.35”) is combined with a social-partner chamber (6.1” × 6.3” × 7.35”) that is separated by a guillotine door with a barrier. This barrier allows for social interaction with olfactory, auditory, and tactile stimulation, but prevents the rats from moving between the test or target chambers. Distinct levers for paired rewards are 6 cm above the grid floors, with discrete light and Sonalert tone cues for social and sucrose-paired levers. The right side of the chamber contains the social-paired lever and the guillotine door for social interaction. The left side of the test chamber is equipped with a sucrose-paired lever, pellet dispenser, pellet receptacle, and an inactive lever. Following behavioral assessment, tray bedding was disposed of, and grates, trays, and the boxes were wiped down with 30% ethanol.

### MedAssociates Social Box Adaptations for Pups

The social operant boxes are a standardized and widely adopted assay intended for adult animals. To safely contain and house pups in the target social chamber, we adopted a series of 3D-printed accessories. We printed a modified barrier with a similar but smaller pattern than the existing barrier, which we affixed to the social door to prevent pups from passing between chambers (Fig. 1A). Similarly, we placed a 3D-printed floor atop the social chamber grate so pups do not slip or get stuck (Fig. 1B). The testing chamber was not altered for this experiment. Following behavioral assessment, all 3D-printed accessories were wiped clean with 30% ethanol.

### Data Collection

The MedPC boxes are wired to a computer, which is used to collect behavioral data using MedPC programs. Subsequently, data was transferred to an Excel file, then analyzed and visualized in GraphPad Prism.

### Subjects

Male and female Long Evans rats were purchased from Charles River Laboratories. Housing conditions involved 20×20×40 cm cages with filter tops (non-ventilated) and *ad libitum* access to food and water. Rats were kept on a 12-hour reverse light/dark cycle, with lights turning off at 10 AM. Males were only used as breeders. Females began lever training for sucrose at postnatal day (PND) 75-100 and were swabbed daily immediately following behavioral assessment using cotton swabs and sterile saline. Vaginal cytology samples were analyzed by hand and scored as being proestrus, estrus, metestrus, or diestrus. Females were pair-housed during sucrose training, then paired for breeding following the final session of sucrose pressing. After a 2-week breeding period, females were single-housed during pregnancy. All female subjects in this experiment were primiparous. The day that pups are born was considered postnatal day (PND) 0. On PND 2, litters were culled to 10 pups (5 males, 5 females – as available) and randomly assigned to press for pups or a female conspecific. Female targets were age-matched and nulliparous. Behavioral studies occurred during the dark phase, an hour after lights out between 12-3 PM.

### Sucrose Self-Administration (SA)

Prior to pregnancy, female rats performed sucrose SA using the Med Associates operant chambers. Critically, this training period familiarized the females to the operant chambers, teaching them that levers are linked to reinforcers. Rats were trained to associate the “sucrose” lever with sucrose reward on 60-minute fixed ratio (FR) schedules. Each sucrose pellet was paired with a 5-second cue light. Animals started training on FR1, where one active lever press results in one sucrose pellet reward. They advanced to FR2 (2 presses for one pellet) and then FR4 (4 presses for one pellet). Training consists of 2 days of each FR session. Given individual differences in learning onset, the animal must earn 5 pellet rewards to qualify for the first day of FR1 training and progress to FR2. If the animal failed to acquire the 5-pellet minimum, they restarted sucrose training. Following FR training, rats were run on one day of progressive ratio (PR) schedule. During PR, the response requirement for each subsequent sucrose pellet delivery increases exponentially until the subject fails to meet the requirement with the following sequence: 2, 4, 6, 9, 12, 15, 20, 25, 32, 40, 50, 62, 77,95, 118, etc. The session expired when an animal took more than 30 minutes to earn a sucrose pellet. The final completed response ratio represents the “breakpoint” value – a numerical quantification of motivation.

### Social Self-Administration

During Social SA, pressing the social lever resulted in a 10-second tone cue, the house light, and the lifting of a guillotine-style door. While the door was raised, the subject and target rat(s) could interact for 30-second. After the interaction, the house light turned off, the guillotine door closed, and a new trial began. Rats were trained in Social SA beginning on PND 11. Dams were randomly assigned to press for their pups (3 males and 3 females, as available) or an unfamiliar female conspecific target. Conspecific targets remained the same throughout testing. Given the virgin sucrose training phase, these females were familiar with lever-reward contingency. We thus truncated the postpartum social self-administration training to 1 day each of FR1, FR2, and FR4. After the 3 days of FR, dams are run on a progressive ratio schedule. This approach allowed us to start training when pups can thermoregulate and do not require additional approaches to stay warm. We shortened the social FR sessions to 30 minutes to avoid extensive maternal separation. A systematic review of maternal studies reveals that separations longer than an hour can be stressful, but those less than an hour are considered brief and consistent with naturally occurring separations in the wild ^48^.

### Behavioral Recording and Video Scoring

During behavioral sessions, animals were recorded in the experimental chambers, using Logitech C920 webcams we affixed to the top of the boxes and OBS Studio software. Webcams were plugged into the same computer used to collect and house MedPC box data. Video recordings were saved locally onto the box computer, then analyzed in Behavioral Observation Research Interactive Software (BORIS). Behavioral scoring was conducted using a standardized ethogram with the following parameters: 1) time spent in the reinforcer zone, defined as orienting toward or engaging with the region of interest (defined below); 2) active lever zone, defined as orienting toward or engaging with the region of interest; 3) self-grooming; and 4) resting. Scoring was performed on the first day of the FR1 schedule and during the final 30 minutes of the social PR session for both sucrose and social tests. Prior to scoring, a region of interest (ROI) was defined surrounding either the social door opening or the active lever. The ROI was created using paint.NET and standardized to extend approximately five bars away from the social door. Due to variability in chamber placement and camera positioning across boxes, a unique ROI was generated for each video. The ROI image was then overlaid onto the corresponding video within BORIS and used to score the area of interest. Behavioral data extracted from BORIS were subsequently exported and analyzed using GraphPad Prism software.

### Statistical Analysis

All presented data are shown as individual data points or group means ± Standard Error of the Mean. Consistent with common practice, datapoints were excluded if they were 1.5 points either above the third quartile or below the first. Statistical significance was analyzed using students paired t-test, Welch’s t-test, ANOVA with Holm-Sidak’s multiple comparison test, when appropriate. P-values less than .05 are considered statistically significant and are visualized with at least a single asterisk. P-values less than .1 are considered trending and depicted with the p-value. P-values greater than .1 are considered insignificant and not shown. Data were visualized and analyzed using GraphPad Prism software.

## RESULTS

### Operant Lever Pressing

Depicted is lever-pressing data over the course of a virgin training period for sucrose (Fig. 3) and a postpartum assessment for social interaction (Fig. 4). For the virgin sucrose pre-training phase, a one-way ANOVA was conducted to examine if pressing differed across training days. There is a significant difference in rewards earned, *F*(5, 24) = 5.26, *p* < .0001, η² = .25, and active lever pressing, *F*(5, 144) = 5.26, *p* < .0001, η² = .33, indicating successful acquisition of the lever-reward paradigm. We do not see an effect of estrous cycle stage on virgin lever-pressing for sucrose, *F*(3, 22) = 1.35, *p =* .28 (Supplemental Fig. 1). Given our focus was on developing this postpartum-pressing protocol, we are underpowered for estrous cycle analysis and our groups are imbalanced. There are conflicting reports on the relevance of estrous cycle stage for reinforcer motivation; it appears that the reward type, duration of acquisition, and experimental parameters are important determining factors^36,49^.

**Figure 3.**
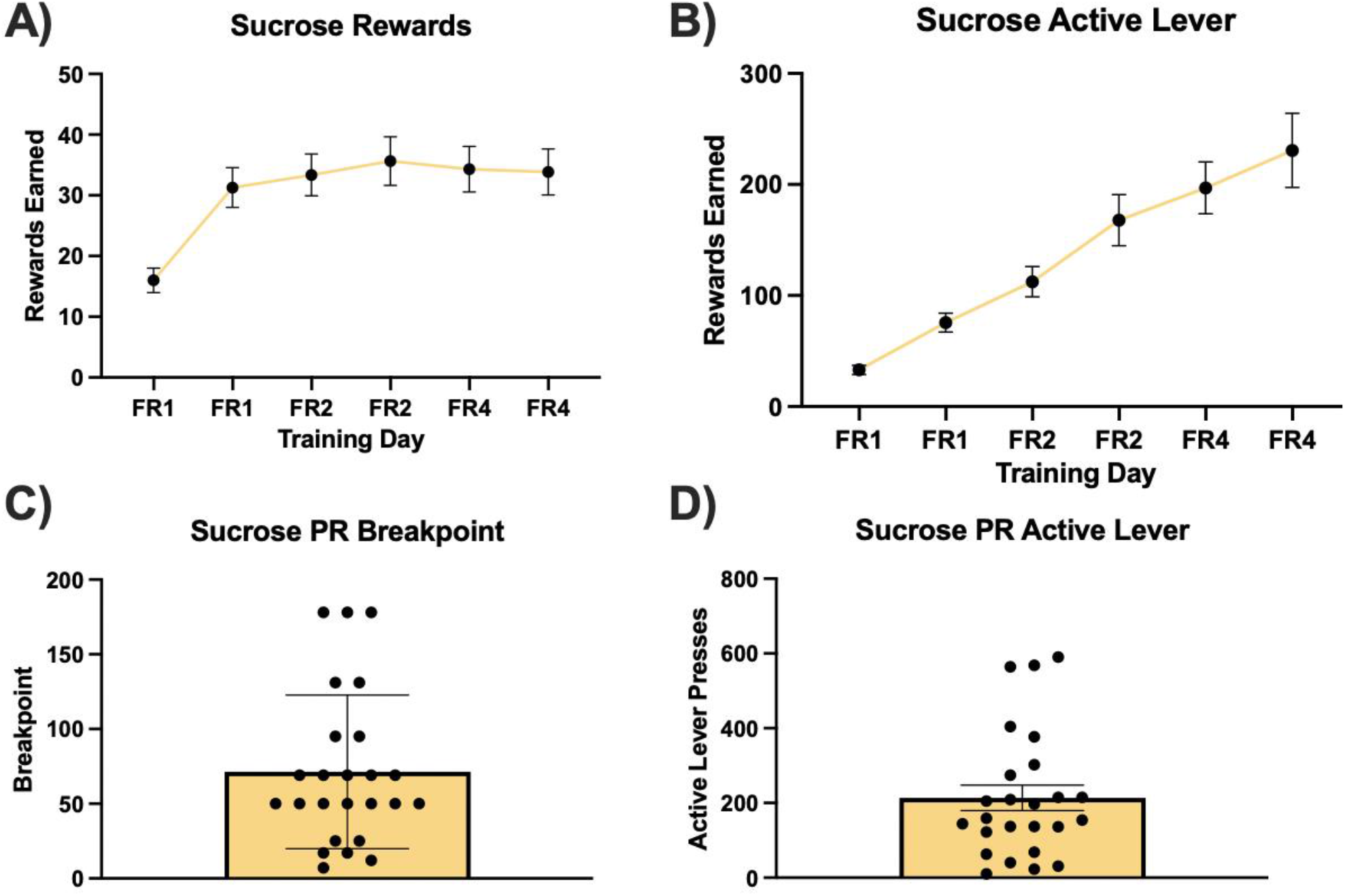
Virgin Sucrose Training. **A)** Female sucrose rewards earned across an FR1, FR2, and FR4 schedule (2 days each), from P11-16. **B)** Female sucrose active lever presses across FR schedule. **C)** Breakpoint of dams pressing for sucrose during progressive ratio (PR) assessment. **D)** Active Lever presses during PR assessment.

**Figure 4.**
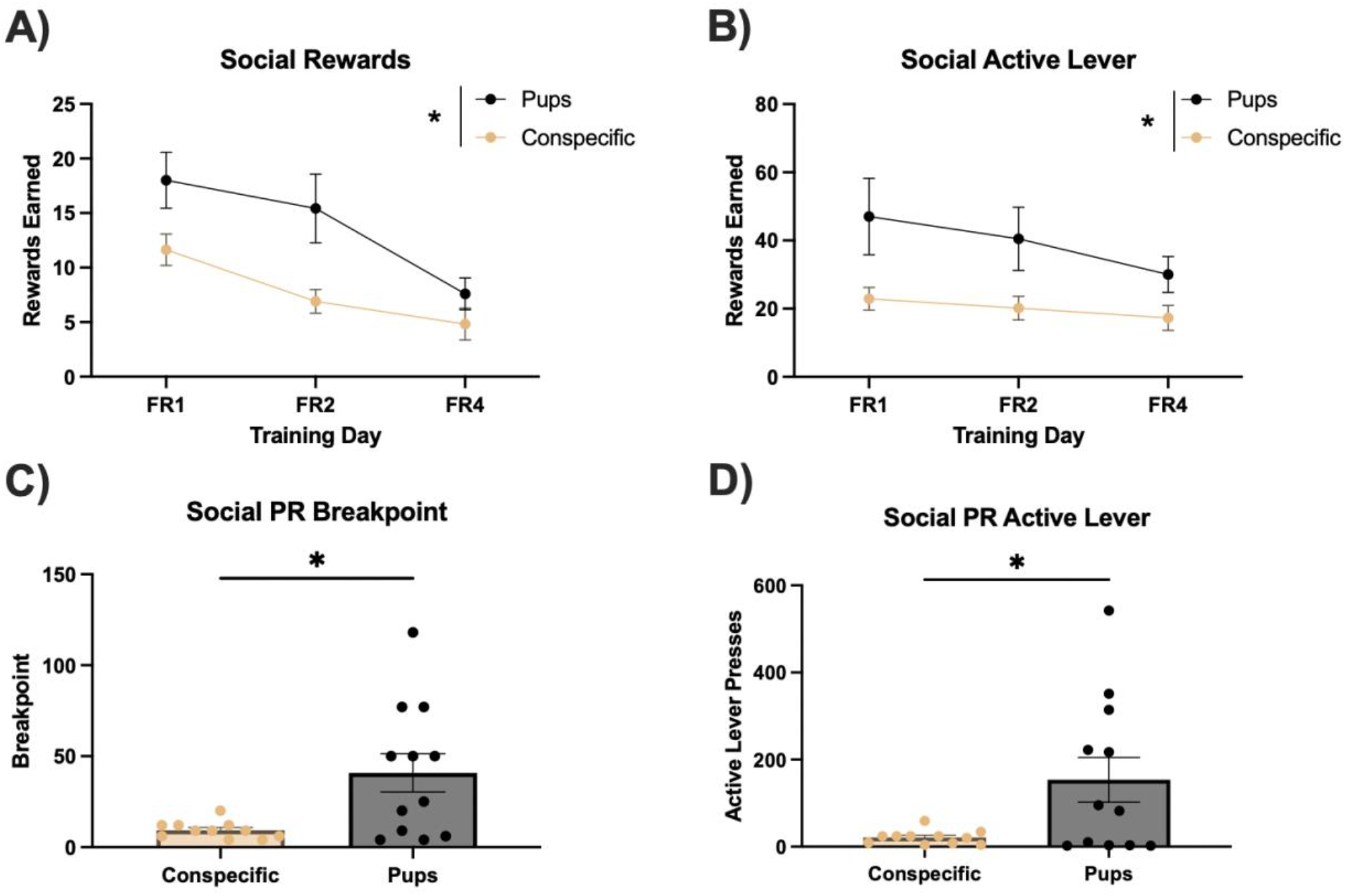
Postpartum Social Assessment. **A)** Dam social rewards earned across FR1, FR2, and FR4 schedule. **B)** Dam social active lever presses across FR schedule. C) Breakpoint of dams pressing for their pups during PR schedule. **D)** Active lever presses during PR schedule. **E)** Correlation between sucrose and social breakpoint. Sucrose and social breakpoints are significantly positively correlated.

For the postpartum social assessment, repeated measures two-way ANOVAs were used to analyze the effect of social target (pup vs. conspecific) and training day on operant outcomes. There are significant or trending main effects of training day on both social reward, *F*(1.94, 40.82) = 20.39, *p* < .0001, and active lever press *F*(1.85, 38.77) = 2.59, *p* = .09. Simple main effects analysis showed that pup-pressers earn significantly higher social rewards (*p =* .03) and press the active lever more (*p =* .03) than dams pressing for a conspecific. There is not a statistically significant interaction between the effect of social target and training day on social reward, *F*(1.94, 40.82) = 2.29, *p* = .12, or active lever press, *F*(1.85, 38.77) = 0.67, *p* = .5. Welch’s t-tests reveal significantly higher PR breakpoints (*p =* .01) and active lever press (*p =* .03) for pups than conspecifics.

### Video Scoring

The data presented show a within-subjects comparison of behavior during the last 30 minutes of sucrose FR1 and social PR (Fig. 5). We chose these timepoints to capture behavioral profiles during PR breakpoint (i.e. peak lever-pressing and motivation) and compare to a motivational baseline for sucrose – a natural reinforcer. During postpartum social PR, dams spend more time in the reinforcer zone (*p <* .01) and the active lever zone (p = .08), compared to when they were virgins pressing for sucrose. Inversely, postpartum dams pressing for their pups show less resting (*p* = .06) and self-grooming (*p =* .1) than the virgin sucrose timepoint. These findings show that postpartum social SA is associated with increased reinforcer-focused but decreased self-directed behaviors.

**Figure 5.**
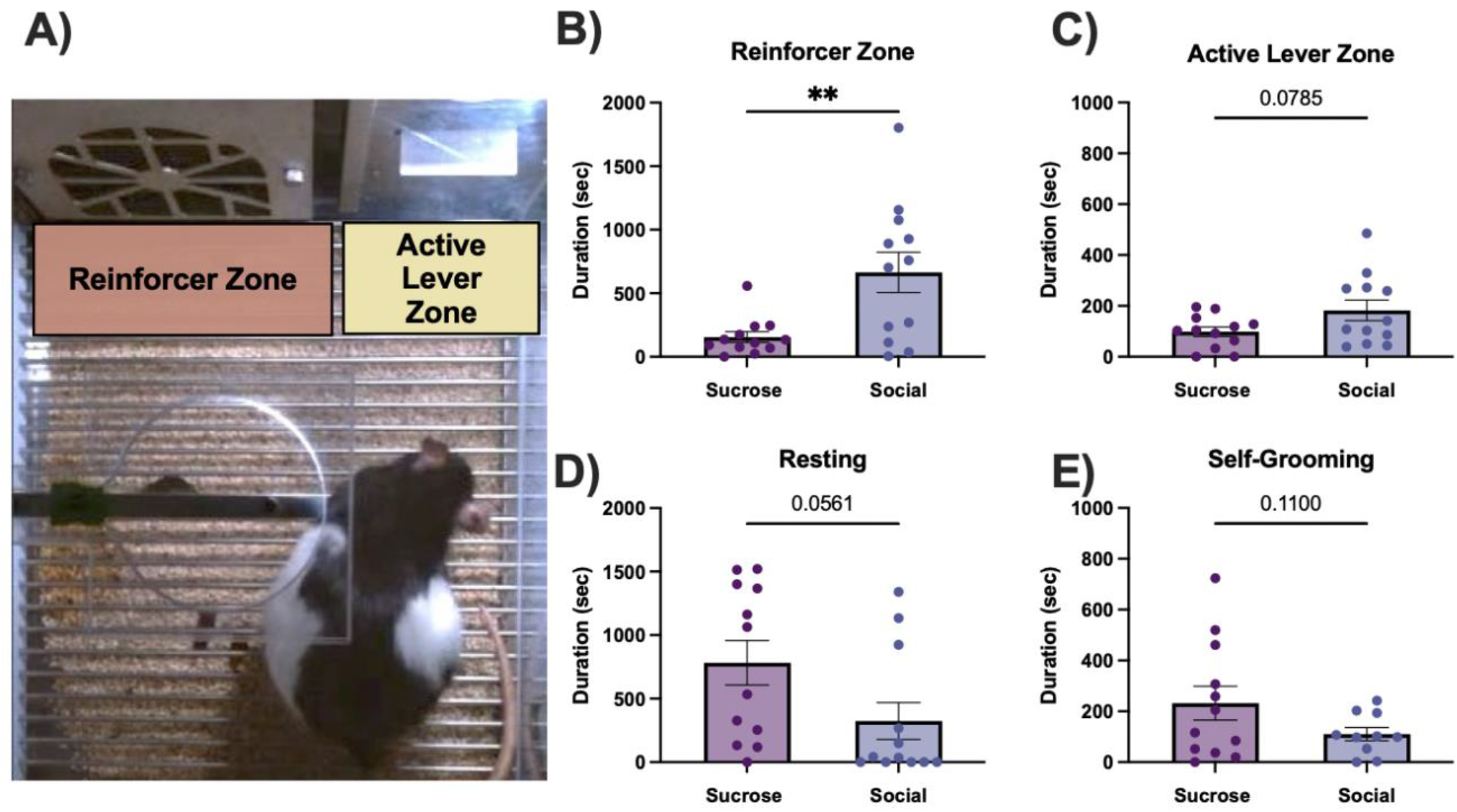
Operant Video Scores. **A)** Example Video Set-Up. Using a within-subjects design, we compared behavior in the operant chamber during sucrose SA pretraining to pup SA. **B)** During postpartum pressing, dams spend more time by the social door for their pups than they do by the sucrose hopper as virgins. C) Dams spend more time in the active lever zone during postpartum social SA than as virgins during sucrose SA. D) Conversely, dams rest less during postpartum pup-pressing than they do during the virgin sucrose pressing. **E)** Dams further show decreased self-grooming during postpartum pup-pressing than during virgin sucrose pressing.

## DISCUSSION

We developed a replicable and accessible approach to investigate postpartum motivation for pups using Med Associates social operant boxes. To our knowledge, this is the first report of postpartum pressing for pups using the Med Associates operant boxes. Given the well-established importance of maternal motivation, this is a major gap in the field. Likely, this gap exists because there are several barriers to testing maternal pressing for pups; the boxes are not meant to house neonatal animals and that pups’ thermoregulation and maternal motivation timelines create a short window for operant training. We addressed these challenges by 3D-printing adaptations to safely contain pups in the social operant chamber and developing a pre-training protocol for postpartum pup pressing.

The described protocol delivers the operant profiles of motivation across female reproductive timepoints and postpartum phenotyping of operant-related behaviors. The key finding of this work is that postpartum dams will press more for their pups than they will for a female conspecific or for sucrose as virgins. Postpartum dams further show a distinct social phenotype of increased reinforcer-directed behavior and decreased self-directed behavior compared to when they press for sucrose as virgins.

This work provides evidence that pups serve as a powerful social reinforcer capable of driving operant responding in postpartum dams. This supports that maternal behavior is governed by reinforcement processes, such as operant conditioning.

### Interpretations and Implications

Here we show that dams will readily engage in an operant lever pressing response for their offspring, supporting that postpartum motivation for pups is a potent social motivator. This aligns with an established body of literature that offspring cues acquire strong motivational salience for postpartum dams – enough to sustain goal-directed, operant responses^33,50,51^. In fact, a prior report shows that adult virgin females have an average breakpoint of 15 for a social interaction with a conspecific^52^. Dams pressing for their pups, however, show an average breakpoint of 41, demonstrating increased social motivation associated with the postpartum period. Further, we identified that this increased sociality is specific to pups, as opposed to general social stimuli, as postpartum dams will barely press for a conspecific (average breakpoint of 8).

This study further implicated that maternal motivation may be conceptualized within the same reinforcement-learning process typically applied to food or drug rewards. In the last decade, there has been an increase in social operant assays, but maternal pup-directed behavior has yet to be examined. Importantly, the present combinatorial operant and video analysis approach allowed us to discern a behavioral phenotype for maternal pup-directed motivation. Dams develop increased motivational salience for offspring in the postpartum period, resulting in increased time spent in reinforcer-associated zones, presumably at the expense of self-directed behaviors.

The paradigm also offers a useful platform for investigating individual differences in maternal motivation. Here we show that postpartum pup-pressing is a highly variable behavior; dams’ social breakpoint for their pups ranges from 4-118. Variability in breakpoint may reflect differences in care quality, stress reactivity, or underlying activation of pup-directed motivational circuits. Importantly, this variability can be leveraged to parse dams into either high or low motivation phenotype animals. There is a plethora of research examining the effect of experience, such as stress, separation, or environment, on quantities of maternal care. The current approach compliments these studies well, as it can provide numerical quantification of motivation that may help us understand why care changes. Further, by thresholding breakpoints and parsing animals into high or low-motivation, this protocol yields a method to test interventions for dysregulated maternal motivation and reverse adverse effects on care.

### Limitations and Future Directions

Several limitations to this approach should be acknowledged. Our selected timepoint was chosen because we were limited by the thermoregulation of the pups, which does not develop until PND11. The reinforcing value of the pups, however, inevitably changes across the postpartum period. Others have demonstrated that, upon pups’ development of executive motor skills, vision, audition, and olfaction around PND16, maternal care decreases to accommodate shifting offspring needs^17,50^. These alterations in care likely correspond with waning motivation. In support of this construct, others have demonstrated that environmental influences, such as resource scarcity stress, induce the most drastic differences in care on PND7 – a strong candidate timepoint for looking at relevant motivational differences^17^. Due to these factors, PND11-16 is a suboptimal window to assess maternal pressing for pups.

The present study does show, however, that dams will display an operant response to pups. Moreover, responding was neither at ceiling nor floor levels, leaving a sufficient dynamic range to examine how environmental conditions modulate maternal motivation. Future studies should innovate methods to safely warm pups, as that is the main barrier to examining alternate timepoints. If pups could remain warm in the operant chamber, the present operant approach could be adapted earlier to identify operant patterns of pressing across the entire postpartum timepoint.

Importantly, this protocol fills an urgent gap in investigating postpartum mechanisms of motivation. There are well-established sex biases in scientific studies, with neuroscience showing the strongest disparity in male vs. female representation^53,54^. While recent years have shown an increase in female inclusion to sex agnostic research questions, there is still a paucity of female-specific or women’s health-focused inquiries^54^. The consequences of these disparities are dire; in the United States, maternal mortality rates have nearly doubled in the last 50 years^55^. In contrast, other high-income countries show either decreases or constancy in maternal mortality outcomes^56^. The relationship between sex bias and disparate health outcomes points to a clear need for more basic research on female-specific mechanisms. This protocol addresses that need by extending standard social operant protocols to postpartum dams and their pups. We intentionally developed this protocol to be accessible, with a low barrier to entry for current social operant box users. With a few minor adaptations, research groups can now incorporate maternal questions into their repertoire of operant approaches – doing their part to bridge the gap in women’s health research.

## Acknowledgements

The authors would like to thank Drs. Marco Venniro and Mat Wimmer for their assistance with protocol design, and Dr. Erin Harris for her help developing the box inserts. This work was supported by the National Institutes of Health (R21 DA059966, R01 DA049837, R01 DA056534, R34 DA061483, R21 DA062844, S10OD032336) and the National Science Foundation (IOS 2313253). The content is solely the responsibility of the authors and does not necessarily represent the official views of the NIH.

## References

1. Orso, R. et al. How early life stress impact maternal care: a systematic review of rodent studies. Frontiers in Behavioral Neuroscience 13, 197 (2019).

2. Champagne, F. A. & Meaney, M. J. Transgenerational effects of social environment on variations in maternal care and behavioral response to novelty. Behavioral neuroscience 121, 1353 (2007).

3. Meaney, M. J. Maternal care, gene expression, and the transmission of individual differences in stress reactivity across generations. Annual review of neuroscience 24, 1161–1192 (2001).

4. Walker, C.-D. et al. Chronic early life stress induced by limited bedding and nesting (LBN) material in rodents: critical considerations of methodology, outcomes and translational potential. Stress 20, 421–448 (2017).

5. Korosi, A. & Baram, T. Z. Plasticity of the stress response early in life: mechanisms and significance. Developmental psychobiology 52, 661–670 (2010).

6. Bath, K., Manzano-Nieves, G. & Goodwill, H. Early life stress accelerates behavioral and neural maturation of the hippocampus in male mice. Hormones and behavior 82, 64–71 (2016).

7. Gallo, M. et al. Limited Bedding and Nesting Induces Maternal Behavior Resembling Both Hypervigilance and Abuse. Frontiers in Behavioral Neuroscience 13, (2019).

8. Guardini, G. et al. Influence of maternal care on behavioural development of domestic dogs (Canis familiaris) living in a home environment. Animals 7, 93 (2017).

9. Sacks, R. M. et al. Childhood maltreatment and BMI trajectory: the mediating role of depression. American journal of preventive medicine 53, 625–633 (2017).

10. Shin, S. H., Miller, D. P. & Teicher, M. H. Exposure to childhood neglect and physical abuse and developmental trajectories of heavy episodic drinking from early adolescence into young adulthood. Drug and alcohol dependence 127, 31–38 (2013).

11. Stern, A. et al. Associations between abuse/neglect and ADHD from childhood to young adulthood: A prospective nationally-representative twin study. Child abuse & neglect 81, 274–285 (2018).

12. Agnew-Blais, J. & Danese, A. Childhood maltreatment and unfavourable clinical outcomes in bipolar disorder: a systematic review and meta-analysis. The Lancet Psychiatry 3, 342–349 (2016).

13. Norman, R. E. et al. The long-term health consequences of child physical abuse, emotional abuse, and neglect: a systematic review and meta-analysis. PLoS medicine 9, e1001349 (2012).

14. Weaver, I. C. et al. Epigenetic programming by maternal behavior. Nature neuroscience 7, 847–854 (2004).

15. Weaver, I. C. et al. Reversal of maternal programming of stress responses in adult offspring through methyl supplementation: altering epigenetic marking later in life. Journal of Neuroscience 25, 11045–11054 (2005).

16. Champagne, F. A. et al. Variations in nucleus accumbens dopamine associated with individual differences in maternal behavior in the rat. Journal of Neuroscience 24, 4113–4123 (2004).

17. Eck, S. R. et al. The effects of early life adversity on growth, maturation, and steroid hormones in male and female rats. European Journal of Neuroscience 52, 2664–2680 (2020).

18. Weaver, I. C. et al. Epigenetic programming by maternal behavior. Nature neuroscience 7, 847–854 (2004).

19. McGowan, P. O. et al. Broad epigenetic signature of maternal care in the brain of adult rats. PloS one 6, e14739 (2011).

20. Meaney, M. J. Epigenetics and the biological definition of gene× environment interactions. Child development 81, 41–79 (2010).

21. Francis, D. & Kuhar, M. Frequency of maternal licking and grooming correlates negatively with vulnerability to cocaine and alcohol use in rats. Pharmacology Biochemistry and Behavior 90, 497–500 (2008).

22. Liu, D. et al. Maternal care, hippocampal glucocorticoid receptors, and hypothalamic-pituitary-adrenal responses to stress. Science 277, 1659–1662 (1997).

23. Champagne, F. A., Francis, D. D., Mar, A. & Meaney, M. J. Variations in maternal care in the rat as a mediating influence for the effects of environment on development. Physiology & behavior 79, 359–371 (2003).

24. Winters, C. et al. Automated procedure to assess pup retrieval in laboratory mice. Scientific reports 12, 1663 (p).

25. Stolzenberg, D. S., Stevens, J. S. & Rissman, E. F. Experience-facilitated improvements in pup retrieval; evidence for an epigenetic effect. Hormones and behavior 62, 128–135 (2012).

26. Zimprich, A. et al. Assessing sociability, social memory, and pup retrieval in mice. Current protocols in mouse biology 7, 287–305 (2017).

27. Beach, F. A. & Jaynes, J. Studies of maternal retrieving in rats I: Recognition of young. Journal of Mammalogy 37, 177–180 (1956).

28. Tzschentke, T. M. Review on CPP: Measuring reward with the conditioned place preference (CPP) paradigm: update of the last decade. Addiction biology 12, 227–462 (2007).

29. Tzschentke, T. M. Measuring reward with the conditioned place preference paradigm: a comprehensive review of drug effects, recent progress and new issues. Progress in neurobiology 56, 613–672 (1998).

30. Pereira, M. & Morrell, J. I. The medial preoptic area is necessary for motivated choice of pup-over cocaine-associated environments by early postpartum rats. Neuroscience 167, 216–231 (2010).

31. Sarkisova, K. Y., Tanaeva, K. & Dobryakova, Y. V. Pup-associated conditioned place preference reaction and maternal care in depressive WAG/Rij rats. Neuroscience and Behavioral Physiology 47, 728–736 (2017).

32. Mattson, B. J., Williams, S. E., Rosenblatt, J. S. & Morrell, J. I. Preferences for cocaine-or pup-associated chambers differentiates otherwise behaviorally identical postpartum maternal rats. Psychopharmacology 167, 1–8 (2003).

33. Seip, K. M. & Morrell, J. I. Increasing the incentive salience of cocaine challenges preference for pup-over cocaine-associated stimuli during early postpartum: place preference and locomotor analyses in the lactating female rat. Psychopharmacology 194, 309–319 (2007).

34. Mattson, B. & Morrell, J. Preference for cocaine-versus pup-associated cues differentially activates neurons expressing either Fos or cocaine-and amphetamine-regulated transcript in lactating, maternal rodents. Neuroscience 135, 315–328 (2005).

35. Wong, S. E. Operant learning theory. HANDBOOK OF 69 (2008).

36. Raymond, J. S., Rehn, S., James, M. H., Everett, N. A. & Bowen, M. T. Sex differences in the social motivation of rats: Insights from social operant conditioning, behavioural economics, and video tracking. Biology of Sex Differences 15, 57 (2024).

37. Skinner, B. F. Operant behavior. *American psychologist* **18**, 503 (1963).

38. Neuringer, A. Operant variability and the evolution of volition. International Journal of Comparative Psychology 27, (2014).

39. Venniro, M. et al. Volitional social interaction prevents drug addiction in rat models. Nature neuroscience 21, 1520–1529 (2018).

40. Neuringer, A. The voluntary operant and the operant nature of volition: Three views. Journal of the Experimental Analysis of Behavior 119, 129–139 (2023).

41. Neuringer, A. & Jensen, G. Operant variability and voluntary action. Psychological review 117, 972 (2010).

42. Lee, A., Clancy, S. & Fleming, A. S. Mother rats bar-press for pups: effects of lesions of the mpoa and limbic sites on maternal behavior and operant responding for pup-reinforcement. Behavioural brain research 100, 15–31 (1999).

43. Hauser, H. & Gandelman, R. Lever pressing for pups: evidence for hormonal influence upon maternal behavior of mice. Hormones and behavior 19, 454–468 (1985).

44. Venniro, M. & Shaham, Y. An operant social self-administration and choice model in rats. Nature protocols 15, 1542–1559 (2020).

45. Farrell, W. J. & Alberts, J. R. Rat behavioral thermoregulation integrates with nonshivering thermogenesis during postnatal development. Behavioral neuroscience 121, 1333 (2007).

46. Kleitman, N. & Satinoff, E. Thermoregulatory behavior in rat pups from birth to weaning. Physiology & Behavior 29, 537–541 (1982).

47. Stone, E. A., Bonnet, K. A. & Hofer, M. A. Survival and development of maternally deprived rats: Role of body temperature. Biopsychosocial Science and Medicine 38, 242–249 (1976).

48. Tractenberg, S. G. et al. An overview of maternal separation effects on behavioural outcomes in mice: evidence from a four-stage methodological systematic review. Neuroscience & Biobehavioral Reviews 68, 489–503 (2016).

49. Richard, J. E., López-Ferreras, L., Anderberg, R. H., Olandersson, K. & Skibicka, K. P. Estradiol is a critical regulator of food-reward behavior. Psychoneuroendocrinology 78, 193–202 (2017).

50. Magnusson, J. E. & Fleming, A. S. Rat pups are reinforcing to the maternal rat: role of sensory cues. Psychobiology 23, 69–75 (1995).

51. Seip, K. M. & Morrell, J. I. Exposure to pups influences the strength of maternal motivation in virgin female rats. Physiology & behavior 95, 599–608 (2008).

52. Williams, A. V. et al. Early resource scarcity alters motivation for natural rewards in a sex-and reinforcer-dependent manner. Psychopharmacology 239, 3929–3937 (2022).

53. Shansky, R. M. & Murphy, A. Z. Considering sex as a biological variable will require a global shift in science culture. Nature neuroscience 24, 457–464 (2021).

54. Beery, A. K. & Zucker, I. Sex bias in neuroscience and biomedical research. Neuroscience & Biobehavioral Reviews 35, 565–572 (2011).

55. Wang, S., Rexrode, K. M., Florio, A. A., Rich-Edwards, J. W. & Chavarro, J. E. Maternal mortality in the United States: trends and opportunities for prevention. Annual review of medicine 74, 199–216 (2023).

56. Zureick-Brown, S. et al. Understanding global trends in maternal mortality. International perspectives on sexual and reproductive health 39, 10–1363 (2013).

