## Supplemental Figure 1 for "Adapting Social Operant Paradigms to Measure Postpartum Maternal Motivation"

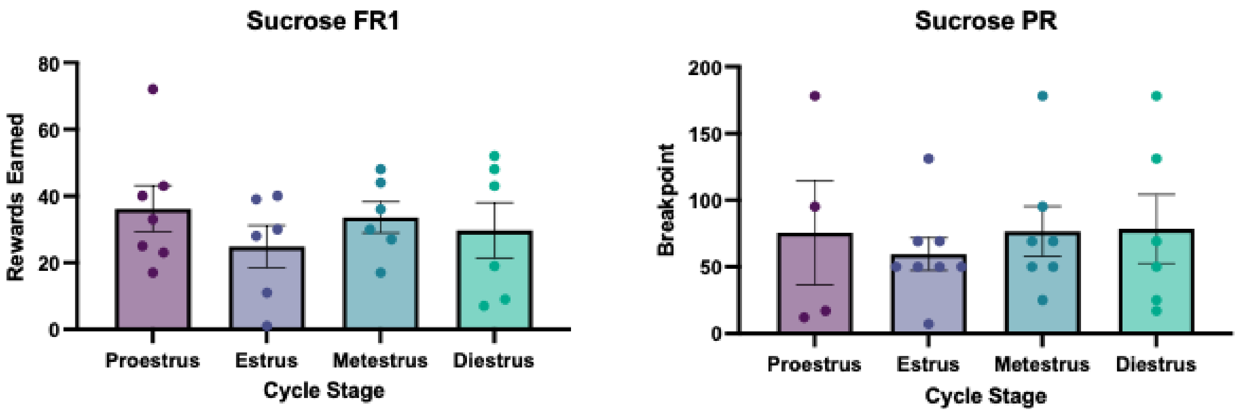


**Supplemental Figure 1. A** )There is no effect of estrous cycle stage on sucrose FR1 rewards earned. **B)** Similarly, estrous cycle does not appear to mitigate sucrose PR breakpoint.
